# Motion direction representation in multivariate electroencephalography activity for smooth pursuit eye movements

**DOI:** 10.1101/519074

**Authors:** Joonyeol Lee, Woojae Jeong, Seolmin Kim, Yee-Joon Kim

**Affiliations:** Center for Neuroscience Imaging Research, Institute for Basic Science (IBS), Suwon 16419, Republic of Korea; Department of Biomedical Engineering, Sungkyunkwan University, Suwon 16419, Republic of Korea; Center for Cognition and Sociality, Institute for Basic Science (IBS), Daejeon 34141, Republic of Korea

**Author notes:** Proofs and correspondence to: Joonyeol Lee, Department of Biomedical Engineering, Sungkyunkwan University, 2066 Seobu-ro, Jangan-gu, Suwon-si, Gyeonggi-do, 16419, Republic of Korea. Voice.

**Keywords:** EEG, inverted encoding model, smooth pursuit eye movement, open-loop, closed-loop, feedback, feedforward, Bayesian inference

## Abstract

Visually-guided smooth pursuit eye movements are composed of initial open-loop and later steady-state periods. Feedforward sensory information dominates the motor behavior during the open-loop pursuit, and a more complex feedback loop regulates the steady-state pursuit. To understand the neural representations of motion direction during open-loop and steady-state smooth pursuits, we recorded electroencephalography (EEG) responses from human observers while they tracked random dot kinematograms as pursuit targets. We estimated population direction tuning curves from multivariate EEG activity using an inverted encoding model. We found significant direction tuning curves as early as 20 ms from motion onset. Direction tuning responses were generalized to later times during the open-loop smooth pursuit, but they became more dynamic during the later steady-state pursuit. The encoding quality of retinal motion direction information estimated from the early direction tuning curves was predictive of trial-by-trial variation in initial pursuit directions. These results suggest that the movement directions of open-loop smooth pursuit are guided by the representation of the retinal motion present in the multivariate EEG activity.

## Introduction

When we navigate through our environments, such as when driving a car, we continuously interact with incoming visual motion information to enact appropriate motor behaviors. This requires our brain to transform sensory motion information into motor commands that actuate the effectors. Smooth pursuit eye movement has been studied to provide insight into how this process occurs in the brain (Lisberger, 2015, 2010). Previous studies have shown that the initiation of the smooth pursuit comprises a feedforward process, known as the ‘open-loop’ period (Krauzlis and Lisberger, 1994; Lisberger and Westbrook, 1985; Tychsen and Lisberger, 1986), followed by subsequent steady-state smooth pursuit where the feedback loop is closed. Neurophysiological recordings in rhesus macaques as an animal model have enabled investigation of the neural mechanisms involved in the feedforward, open-loop pursuit. Most of the behavioral variation in visually guided smooth pursuit at the end of the open-loop period can be explained by sensory estimation errors (Osborne et al., 2005). Single cell recording studies showed that these behavioral errors can be explained by the correlated neural variation in macaque visual area MT (Hohl et al., 2013; Lee et al., 2016; Lee and Lisberger, 2013), without any additional variation being added in downstream motor areas (Medina and Lisberger, 2007; Schoppik et al., 2008).

In contrast, we have limited understanding of how neural activity in the human brain represents information processes in smooth pursuit eye movements. Functional magnetic resonance imaging studies revealed brain areas that are important for sensorimotor transmission (Berman et al., 1999; Burke and Barnes, 2008; Nagel et al., 2008, 2006; O’Driscoll et al., 2000; Petit et al., 1997; Petit and Haxby, 1999; Rosano et al., 2002; Tanabe et al., 2002). However, due to the poor temporal resolution of this method, it is difficult to dissect the differences in the temporal dynamics of neural activity during open-loop and closed-loop periods. In several studies, electroencephalography (EEG) responses were measured during smooth pursuit eye movements, taking advantage of the high temporal resolution of EEG. However, most of these studies focus on the neural correlates of cognitive factors during smooth pursuit (Chen et al., 2017a, 2017b). Studies investigating how basic features of motion information (speed, direction, and timing) are represented in the human brain during sensorimotor behaviors are scarce. This lack of knowledge is partly due to methodological limitations in linking large-scale population activity measured in the human brain to lower dimension features. Recent advances in the multivariate analysis of neural data have shed light on elucidating this possibility (Naselaris et al., 2011). In particular, the use of encoding models have proven fruitful in examining population responses for lower level features, such as color, orientation, speed, and direction of motion (Brouwer and Heeger, 2013, 2011, 2009; Chong et al., 2015; Garcia et al., 2013; Kok et al., 2013; Vintch and Gardner, 2014; Wolff et al., 2017)

In this study, we investigated how directional information of sensory and pursuit motion during a smooth pursuit eye movement task is represented in multivariate EEG activity of human participants. Instead of focusing on individual channel responses, we observed the motion information present in the pattern of EEG activity across all recording channels using an inverted encoding model (Brouwer and Heeger, 2009). We found robust direction tuning responses from approximately 80 ms before the initiation of smooth pursuit to the steady-state period. Investigation of temporal dynamics revealed distinctive patterns of direction tuning curves in open-loop and closed-loop period pursuits. Further, encoding quality of motion direction representation was predictive of the size of pursuit direction errors and variability in pursuit directions during the open-loop period. This relationship can be easily explained if we understand the initiation of smooth pursuit as a result of Bayesian inference.

## Materials and Methods

### Subjects

EEG data and smooth pursuit eye movement data were collected from 21 human participants. Before each EEG recording session, we conducted behavioral training for 1 hour to screen subjects whose performance in the smooth pursuit eye movement task was poor (defined as too many saccades at the initiation of smooth pursuit) and to train human participants on the task. EEG recordings were scheduled and conducted on different days. We discarded subjects from further analysis according to behavioral and neurophysiological criteria listed below, after initial data analysis. Of 21 participants, five subjects were excluded from further analysis due to frequent saccadic eye movements during fixation and pursuit (if more than 30% of the trials were discarded due to saccadic eye movements, participants were excluded from further analysis). One participant was excluded from analysis because of poor EEG recording quality (more than 50% of ICs were removed after running ADJUST algorithm to isolate artifactual components, see below).

All participants had normal or corrected-to-normal vision. We obtained written consent from all participants before each experiment, and all experimental protocols were approved by the Institutional Review Board at Sungkyunkwan University.

### Stimuli and task design

The task was a simple smooth pursuit eye movement task, modified from a step-ramp pursuit task after Rashbass (Rashbass, 1961). Figure 1 illustrates the task design. We displayed the visual stimuli on a gamma corrected 20-inch CRT monitor (Hewlett Packard p1230) that was positioned 60 cm from the participants. It covered 36.9° by 28.1° of the visual field. The spatial resolution of the monitor was 1600 by 1200 pixels. The vertical refresh rate was 85 Hz. All visual stimuli were presented on a gray background in a grayscale that covered a luminance range from 0 to 72.5 cd/m².

**Figure 1:**
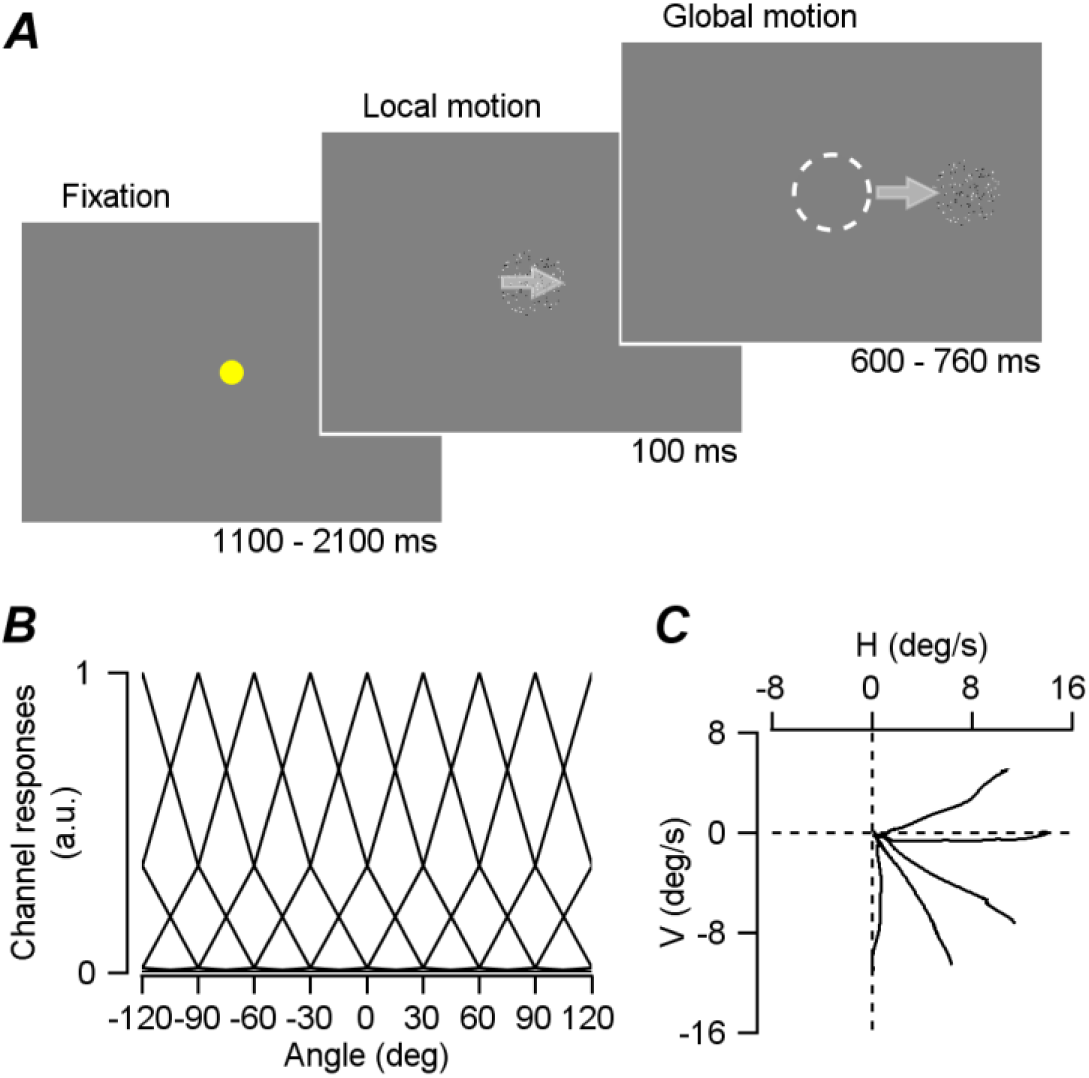
Task design and forward encoding model ***A.*** A simple schematic of the smooth pursuit eye movement task. After the random duration of fixation (1100 to 2100 ms), a brief pattern motion appears (100 ms) at a speed of 16 /s and one of the five directions is randomly selected. Window motion followed in the same speed and direction. ***B.*** Channel responses used in the inverted encoding model. We used 11 direction channels that encompassed −180 to 120 in 30 steps. Half-cosine function raised to the 15th power was used in modeling the channel responses. ***C.*** Mean eye velocity traces of one subject in the open-loop period (from 100 to 220 ms from motion onset) of smooth pursuit.

We used random dot kinematograms as the pursuit target. The dot patch comprised 128 spots inside a 4.5°-diameter circular patch. Nominal contrast of the dot patch was 100% because we randomly mixed equal numbers of minimum luminance dots (black) and maximum luminance dots (white) in the patch (Yang et al., 2012). Therefore, the mean luminance of the patch was the same as that of a gray background. All experiments were conducted in a reasonably dark room, with the display monitor as the main source of the illumination.

Each trial started with a yellow fixation spot presented at the center of the display. Subjects had to fixate on the fixation spot within a 2° square invisible window, for a random duration between 1100 and 2100 ms. Immediately after the extinction of the fixation point, random dot kinematograms appeared at the center of the display and started moving at 16°/s in one of the five predetermined directions (0°, ±30°, 270°, and 300°, Figure 1C). For the first 100 ms, the invisible circular aperture of the random dot patch was stationary, but all dots inside the aperture moved in 100% coherence. Then, all dots and the aperture moved together in the same speed and direction for a random duration between 600 ms and 790 ms. Individual trials were considered successful only if subjects tracked the random dot kinematograms within a 5° invisible square window around the center of the dot patch until the target stopped moving and disappeared from the screen. Only successful trials were saved for further analysis. There were 1000 ms pauses between each trial. Every 80 trials, subjects were given a 1-minute break.

### Data acquisition

Horizontal and vertical eye positions of the right eye of each participant were monitored and saved through an infrared eye tracker (EyeLink 1000 Plus, SR research), with a sampling rate of 1 kHz. Control of visual stimuli and acquisition of eye position signal were performed using a real-time data acquisition program (MAESTRO) used in previous studies (https://site.google.com/a/srscicomp.com/maestro/). To guarantee the timing of visual stimuli, we used a custom-built photodiode circuit. Every EEG and behavioral data were aligned to the photodiode markers. We used a 64-channel EEG amplifier (BrainAmp, Brain Products, GmbH) and active electrodes (actiCAP, Brain Products, GmbH) mounted in an elastic cap for recording EEG signals from human observers. The impedances of the electrodes were kept under 5 kΩ during the recordings. The sampling rate of EEG data collection was 5 kHz. All behavior control-related information was transmitted to the EEG data collection computer online through a custom-built hardware interface and digital IO for subsequent synchronization.

### Preprocessing of EEG data

Before we analyzed the recorded EEG data, we removed artifactual components that originated from sudden changes in electrode impedance and eye movements through a series of preprocessing procedures. All preprocessing and further analysis were performed using subroutines of EEGLab (Delorme and Makeig, 2004) and FieldTrip (Oostenveld et al., 2011) Matlab (MathWorks, Inc.) toolboxes. First, we down-sampled the EEG data from 5 kHz to 1 kHz and applied highpass filter with 1 Hz cutoff frequency for removing slow drifts (Winkler et al., 2015) and a lowpass filter with cutoff frequency of 200 Hz. We used Artifact Subspace Reconstruction (ASR) routine (Mullen et al., 2015) to remove noisy channels, and re-referenced all the data using the average as the reference (Bigdely-Shamlo et al., 2015). We removed line noise (60, 120, and 180 Hz) using the *cleanline* EEGLab plugin (Mullen, 2012). Then, the artifactual response components induced by eye movements and blinks were identified via independent component analysis (ICA) (Makeig et al., 1996). Finally, we determined ICs that were classified as artifacts using *ADJUST* EEGLab plugin (Mognon et al., 2011), and removed them from the data.

### Inverted encoding model

To extract the direction information of visual motion and smooth pursuit eye movements from spatially distributed EEG activity, we used the inverted encoding model (Brouwer and Heeger, 2009). We used 11 hypothetical direction channels from 120° to −180° in 30° steps, each with an idealized direction tuning curve (half-cosine basis function raised to the 15th power). For this analysis, we first smoothed the EEG responses recorded in each electrode using 30 ms rectangular time window, with 5 ms step size, from - 100 ms to 600 ms relative to visual motion onset (for the population direction tuning curves during the smooth pursuit eye movements), or with 5 ms step size, from −100 ms to 400 ms relative to visual motion onset (for the trial-by-trial correlation analysis). EEG activity was baseline-corrected by subtracting average EEG activity between −100 and −50 ms from the motion onset in individual trials. We normalized each subject’s EEG data by subtracting mean EEG responses across all channels, time points, and trials from the smoothed EEG data and by dividing them by standard deviation across all channels, time points, and trials. Then, at each time point, we estimated the weight matrix for the eleven direction channels using a generalized linear model.

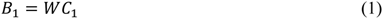

We used the leave-one-condition-out method to prevent overfitting in the estimation. We used the EEG data from all other direction conditions (four directions) excluding the current direction in the weight matrix estimation. *B*_1_ represents EEG data collected from *m* recording electrodes and *n*_1_ trials (*m* × *n*_1_) from four direction conditions, *C*_1_ represents the hypothetical direction channels (*c* × *n*_1_, where *c* is the number of the direction channels), and *W* represents the weight matrix (*m* × *c*).

Estimation of the weight matrix was performed through the ordinary least-square procedure following the equation below.

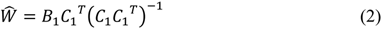

Then, we estimated the directional channel responses using the estimated weight matrix from equation (2).

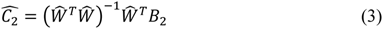

Where 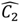 is the estimated channel response for a given direction condition (*c* × *n*_2_), *B*_2_ is EEG data for the direction condition (*m* × *n*_2_). We repeated this calculation across all time points. Since we collected EEG data for five direction conditions without covering the full twelve directions (due to practical reasons), we corrected channel response estimation using a bootstrap procedure. First, for a given direction condition, we estimated the weight matrix using four other direction conditions using equations (1) and (2). Then, we shuffled the trials from all direction conditions and randomly selected samples whose size matched the sample size of the current direction condition. We repeated this 1000 times and made the ‘null’ distribution for the channel response estimates. We converted the channel response estimates into standard scores by subtracting the mean and dividing by the standard deviation of the null distribution and used them in subsequent analyses.

When we performed the cross-temporal generalization analysis, we estimated the weight matrix using EEG data at each time point and applied the weight for the estimations of channel responses across all time points. Leave-one-condition out method and bootstrap procedure were used. Encoding accuracy, used in temporal generalization and group comparison analysis, was estimated from the sharpness of the z-score of direction tuning responses (Myers et al., 2015). We used the slope of linear regression between direction and channel responses as the index of sharpness. We averaged the responses of the leftward four direction channels in ascending order (−120, −90 −60 −30) and responses of the rightward four directions channels in descending order (120, 90, 60, 30), relative to nominal center direction. Using the four average channel responses and the center direction channel response, we performed linear regression analysis and estimated the slope. When we evaluated the accuracy of encoding from individual trials, we used how well a von Mises distribution explained the direction tuning responses as an index. Instead of using the tuning curve slope, we obtained this measure for a more robust index of tuning quality from single trials. We fitted the direction tuning curves in individual trials to the von Mises distribution using the least square procedure.

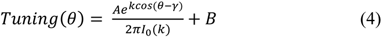

In this equation, *A* is scaling factor, *I*_0_(*k*) is the zeroth order Bessel function, θ is the direction of motion, *γ* is the preferred direction, and *B* is offset. The amount of variance of the direction tuning responses that is explained by a von Mises distribution was used as the index for encoding accuracy with values from zero to one. One means that all of the tuning response variation is explained by the von Mises distribution.

### Analysis of smooth pursuit eye movements

To quantify the initiation of smooth pursuit eye movements, we obtained horizontal and vertical velocity traces by differentiating eye position data that were obtained from the infrared eye tracking system. Before the differentiation, we discarded any high-frequency components by applying 6 pole low-pass Butterworth filter, with a cut-off frequency of 25 Hz using a subroutine of FieldTrip Matlab package. Then, we screened all the velocity traces in individual trials visually and discarded trials that contained saccadic eye movements in a time window between −100 and 250 ms from visual motion onset (Figure 2B). This practice enabled us to analyze the open-loop period of the smooth pursuit (Lisberger et al., 1987; Lisberger and Westbrook, 1985) and prevent abrupt eye movements from affecting EEG responses in the time of interest. To quantify the pursuit direction during the initiation of pursuit, we averaged horizontal and vertical eye velocities using a 30 ms time window, from 0 to 200 ms relative to mean pursuit latency of each participant and took arctan of the two components. Latencies of smooth pursuit for individual participants were estimated by a statistical criterion. On each pursuit condition, we calculated mean eye speeds as a function of time and compared the speed at each time point with baseline eye speed (−50 ms to 20 ms from motion onset). If the average eye speed exceeded 0.3 standard deviations from mean baseline eye speed for 50 ms, then we took the first time point that exceeded the criterion as the latency. This measure usually matched the pursuit latency that was manually selected through visual inspection.

**Figure 2:**
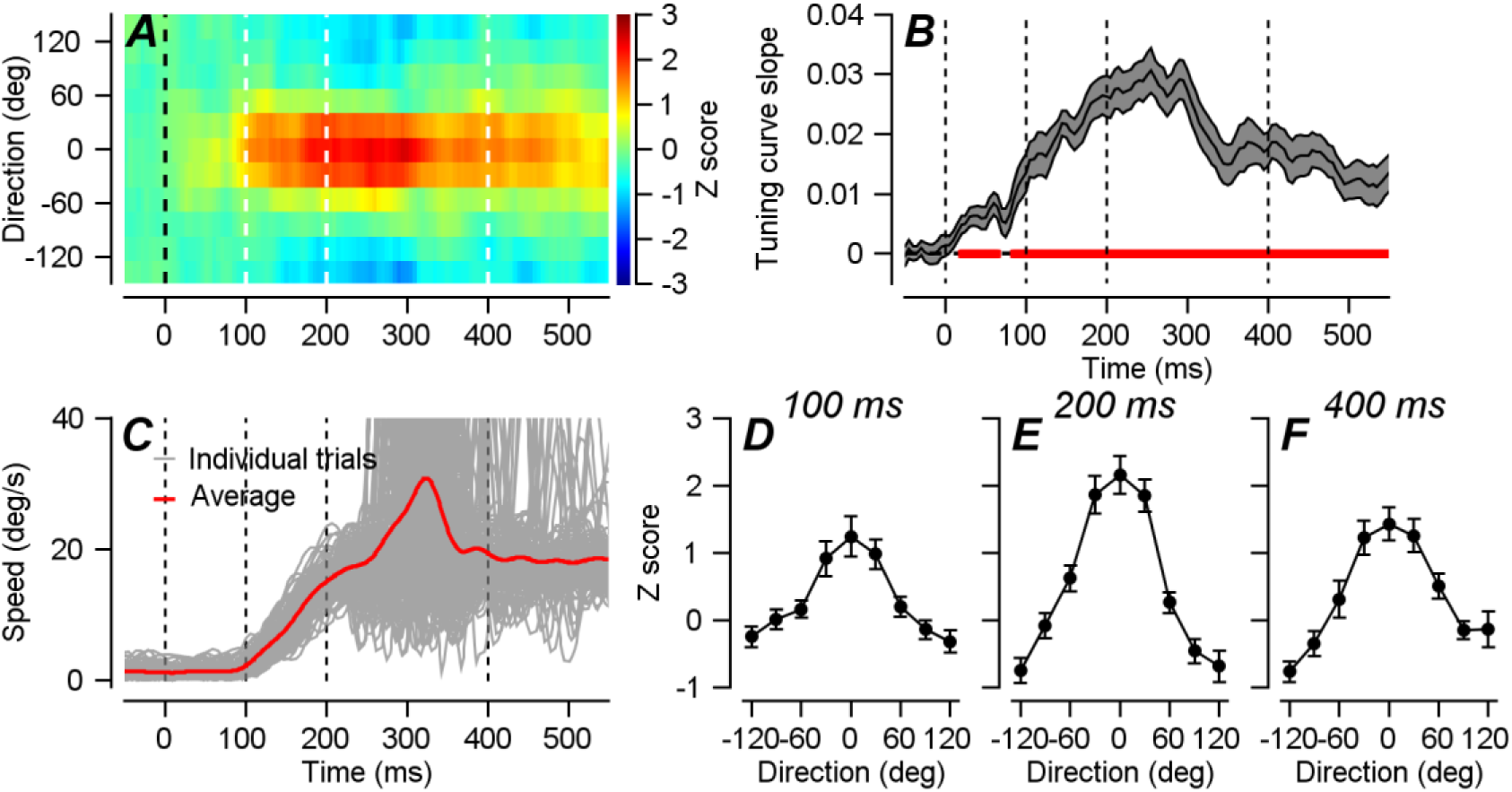
Direction channel responses estimated from EEG activity using the inverted encoding model. ***A.*** Direction tuning responses estimated from 64-channel EEG activity. Visual motion appeared at time zero, and color shows the population direction tuning curves that are converted into standard score through permutation analysis. ***B.*** Encoding accuracies estimated from the slope of the direction tuning curves (see *Methods* for details). The red colored line shows the time points where encoding accuracy is significantly different from zero (cluster-based permutation test, p < 0.01). Gray areas show standard errors. ***C.*** Example traces of smooth pursuit eye movements. Trials that included saccadic eye movements in the time window between −100 and 250 ms from motion onset were removed from further analysis. ***D-F.*** Population direction tuning curves estimated at 100 (***D***), 200 (***E***), and 400 (***F***) ms from motion onset. Error bars show standard error.

When we compared the encoding accuracy with pursuit direction in individual trials, we performed another screening process to remove trials with poor pursuit eye movements. We used a pursuit decomposition procedure that was previously developed (Lee and Lisberger, 2013). If the fitted eye traces in individual trials explained less than 50% of the original eye velocity trace, we did not include them in further analysis, which effectively removed trials with poor pursuit initiation.

## Results

Human participants were instructed to perform a simple smooth pursuit eye movement task where random dot kinematograms were the pursuit target (Yang et al., 2012). We recorded eye position using infrared video camera and EEG activity using 64 channel active electrodes from 15 human observers (12 male). We observed that 1) sensory motion direction information in smooth pursuit could be decoded from multivariate EEG activity as early as 20 ms from motion onset, 2) EEG representation of visual motion direction was generalized to the motion direction representation in EEG during open-loop smooth pursuit but not during steady-state period, and 3) encoding accuracy of EEG activity for visual motion direction was predictive of the size of pursuit direction error and direction variability in the open-loop period.

### Encoding of visual and pursuit motion direction

To extract the direction information of visual motion and smooth pursuit, from multivariate EEG activity, we used the inverted encoding model that was developed by Brouwer and Heeger (Brouwer and Heeger, 2009). We modeled the hypothetical direction channels with direction tuning curve using half-cosine basis function and transformed EEG activity in 64-channels sensor space to responses in the space of 11 hypothetical direction channels (see Methods and Figure 1B). Figure 2A shows the average population direction tuning curve across the 15 participants as a function of time. The visual motion direction information encoded in EEG signals gradually emerged immediately following motion onset. Encoding accuracy, estimated from the slope of the direction tuning curve (see *Methods*, Myers et al., 2015), was significantly different from zero (two-sided cluster-based permutation test, 5000 permutations, p < 0.05) as early as 20 ms from motion onset (Figure 2E). For this analysis and subsequent analyses, we discarded trials with saccadic eye movements in the time window between −100 ms and 250 ms from motion onset (Figure 2B), to ensure that EEG activity used for the analysis was free from the effect of abrupt eye movements (in addition to the removal of eye movement and blink artifacts through the combination of ICA and ADJUST, see Methods for detail). Given that initiation of smooth pursuit started at around 100 ms (Figure 1B) from motion onset, direction tuning responses from 20 ms to around 120 ms arose from visual motion (Figure 2A, 2C). Subsequent direction tuning responses were stronger than the pure visual motion direction responses (Figure 2D). The stronger direction responses later on may be due to the combined EEG representations of sensory and motor directions. The direction tuning responses were quickly attenuated from 300 ms after motion onset. The reduction in direction responses is likely due to the lack of retinal motion during the steady-state pursuit. Because the feedback loop is closed during the steady-state pursuit, motion direction representation encoded in EEG activity is expected to be temporally more dynamic (see below).

### Cross-temporal generalization of encoding direction information

Next, we used cross-temporal generalization (CTG) analysis to understand the underlying neural dynamics of the direction tuning responses that emerged as early as 20 ms from motion onset. Transformation weight from electrode sensor space to direction channel space was estimated at each time point and applied to all other time points to predict direction tuning responses. Figure 3A shows the cross-temporal generalization map that was composed of the encoding accuracy. The pattern of cross-temporal generalization may provide clues about whether the population direction tuning curves were dynamically changing or maintained across time (King and Dehaene, 2014). If the initial direction tuning curves were qualitatively different from the later tuning curves, EEG activity related to motion direction earlier in time cannot be generalized to EEG activity later in time; therefore, it would show a diagonal pattern. We found that the population direction tuning curves estimated from early sensory EEG activity could be generalized to the tuning curves of later times during the open-loop smooth pursuit (Figure 3A, cluster-based permutation test, 5000 permutations, p = 0.0002). For example, when we estimated the transformation weight matrix at 50 ms from motion onset and applied it to the EEG activity at 200 ms for obtaining the direction tuning curve, the tuning curve was quite robust. The tuning curve was even sharper than the direction tuning curve at 50 ms (Figures 3C and D). This suggests that EEG representation of direction responses are similar across different times, but the signal-to-noise ratio of the underlying neural activity for motion direction becomes greater at a later time (Liu et al., 2018). Further, the lower parts of the off-diagonal encoding accuracies were slightly higher than the upper parts. The asymmetry suggests that the population direction tuning responses at earlier times is predictive of later direction responses, but not vice versa. The gradual increase in signal-to-noise ratio and subtle asymmetry can be well explained by the ramping type of underlying neural responses (King and Dehaene, 2014). In sum, we conclude that the motion direction information represented in multivariate EEG activity is mostly determined by the initial responses to sensory motion and maintained throughout the open-loop period.

**Figure 3:**
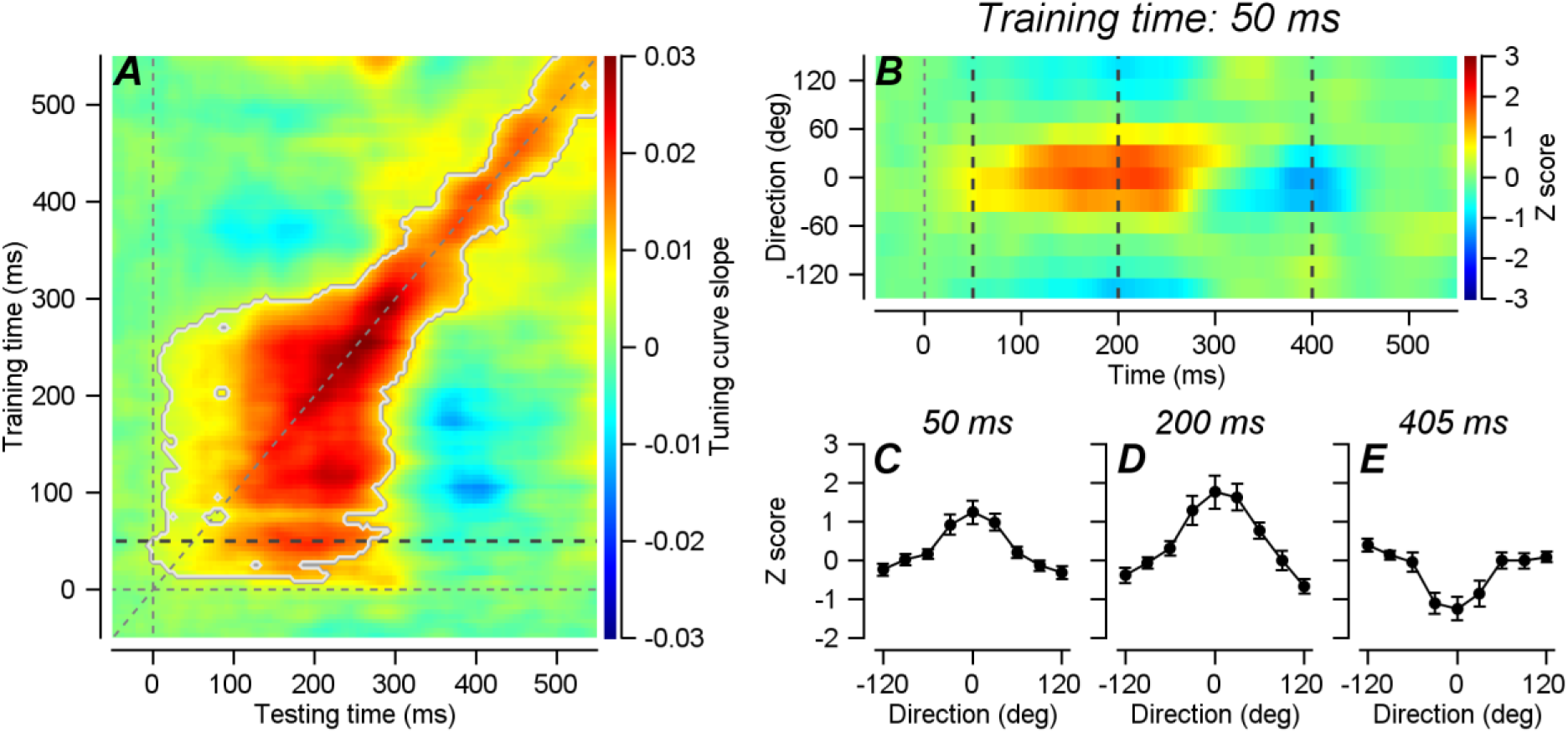
***A.*** Cross-temporal generalization (CTG) of the encoding accuracy of direction tuning responses. The x-axis shows testing time, and Y-axis shows training time. Encoding accuracy was estimated from the sharpness of the tuning response through linear regression. Gray contour shows significant times where encoding accuracies are significantly different from zero (cluster-based permutation test, 5000 permutations, p < 0.001). ***B.*** Direction tuning curves when the transformation weight was trained by EEG activity at 50 ms from motion onset. Color shows tuning curves transformed into standard score through permutation analysis. Tuning curves are sharper at later times, suggesting that EEG activity patterns related to the given motion direction appearing at 50 ms becomes stronger and evolves during the open-loop pursuit. See text for details. ***C-E.*** Population direction tuning curves measured at 50 (***C***), 200 (***D***), and 400 (***E***) ms from motion onset. Error bars show standard error.

The pattern of direction tuning response dynamics appears to be different in the later steady-state pursuit. Significant encoding only appears in the diagonal of the cross-temporal generalization matrix (from 300 ms to 550 ms). At steady state, a moving visual target would be stabilized at the fovea by the closed-loop feedback circuit. Therefore, directions of retinal motion and corrective eye movements would dynamically change, and multiple direction-specific activity patterns may have been triggered in sequence (Figures 3A, E, King and Dehaene, 2014).

### Encoding the accuracy of visual motion direction is predictive of the size of the behavioral error

Through the cross-temporal generalization analysis, we have shown that the population direction tuning responses estimated from EEG activity during the open-loop smooth pursuit can be generalized across time. This result demonstrates that neural *sensory* motion representation guides and dominates the neural representation of *motor* responses in feedforward sensorimotor behavior. As the next step, we tested if the accuracy of the neural representation for motion direction is predictive of the accuracy of motor behavior.

We estimated the encoding accuracy of EEG multivariate activity in a central smooth pursuit direction condition (−30°, see *Methods*) at every trial and time, and compared them with the size of behavioral direction error. In this case, we used how much variation in the estimated direction tuning responses was explained by a hypothetical tuning curve, as an index of encoding quality (see *Methods*). We used this value to obtain more robust indexes from a single trial. We estimated the encoding quality of motion direction in five distinct time windows (25 – 75 ms, 50 – 100 ms, 75 – 125 ms, 100 – 150 ms, and 125 – 175 ms from visual motion onset). Figure 4A shows average direction tuning responses obtained at the time window of 50 to 100 ms from motion onset. Tuning curves of high and low encoding quality are plotted in black and gray, respectively. Figure 4B shows the tuning responses as a function of time. The upper and lower figures indicate when tuning quality is high and low, respectively (see below for how we divided trials). Even when we used how well the tuning responses were fitted by a von Mises distribution (see Methods), tuning curves in high accuracy trials were clearly better than curves in low accuracy trials (Figure 4A, B).

**Figure 4:**
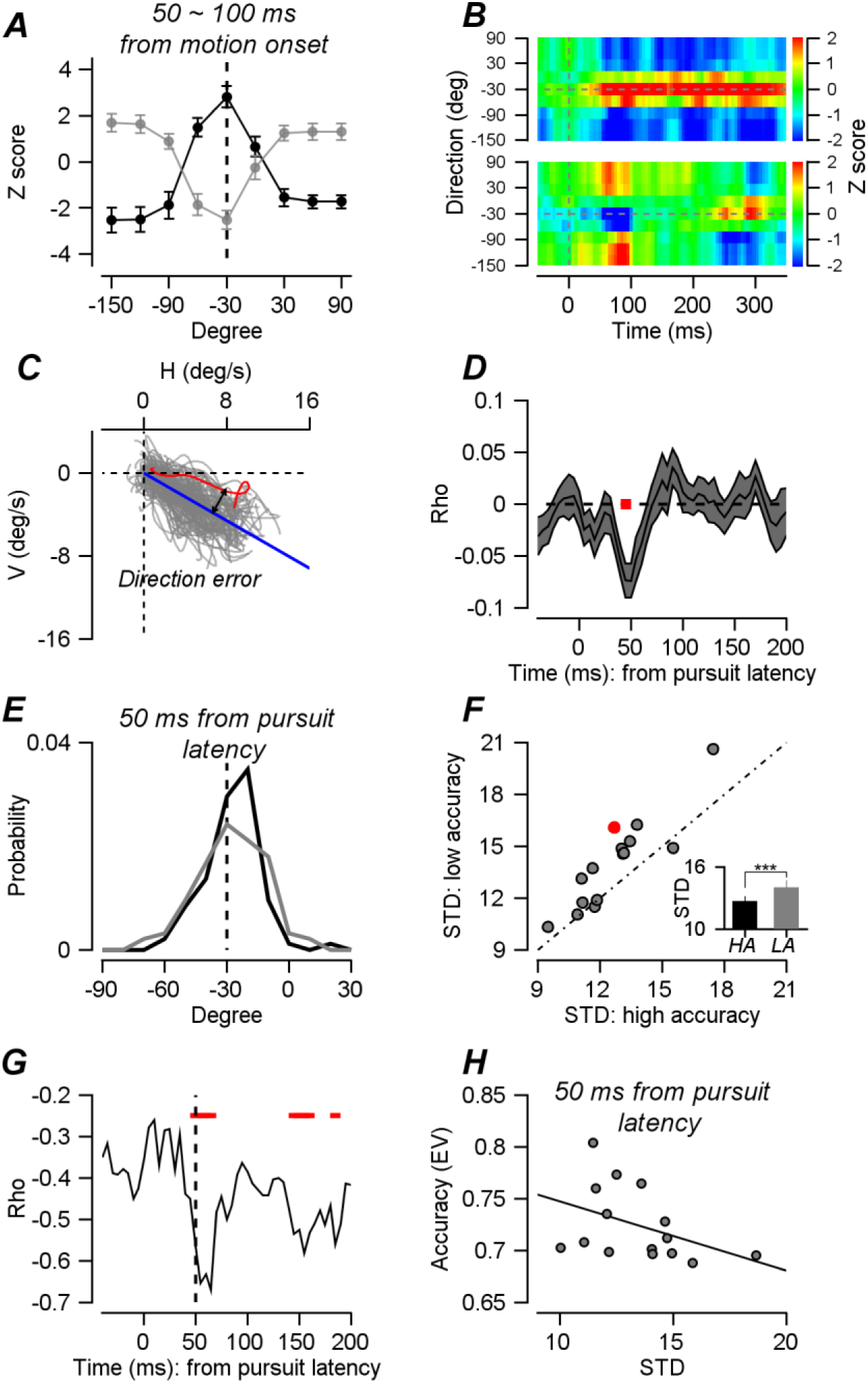
Neural correlates of behavioral direction errors and variation. ***A.*** Direction tuning curves estimated at 50 to 100 ms from motion onset using the inverted encoding model. The black line shows the average tuning curve with high encoding accuracy trials; the gray line shows the average tuning curve with low encoding accuracy trials. Error bars indicate standard error. ***B.*** The tuning curves presented as a function of time. The upper figure is when encoding accuracy at 50 to 100 ms is high, and the lower figure is when encoding accuracy is low. ***C.*** Direction errors were calculated by the difference between the mean pursuit direction and individual pursuit directions of open-loop pursuit velocity. ***D***. Trial-by-trial correlations between the size of direction error and encoding accuracy averaged across the 15 subjects. The dark gray-colored band shows standard error, and the red bar shows the time points where the correlations are statistically significant (from 40 ms to 50 ms relative to pursuit latency, cluster-based permutation test, 5000 permutations, p = 0.007). ***E.*** Example distributions of pursuit direction measured at 50 ms from pursuit latency. Pursuit trials are divided by the encoding accuracies estimated at 50 to 100 ms from motion onset. The black line indicates trials with high encoding accuracy, and the gray line indicates trials with low encoding accuracy. ***F.*** Scatter plot of 15 participants' pursuit direction standard deviation (STD) comparing high encoding accuracy trials with low encoding accuracy trials. The red dot is the example participant’s data. The inset shows the summary of the effect of encoding accuracy on the variation of pursuit direction across the 15 participants. ***G.*** Correlations between average encoding accuracies and STDs of pursuit direction of the 15 participants. STDs of pursuit directions are measured from −50 ms to 200 ms relative to pursuit latency. Red colored bars show time points where the correlations are statistically significant (Spearman's rho, p < 0.05). ***H.*** Scatter plot of the average encoding accuracies and pursuit direction STDs measured at 50 ms from pursuit latency. The linear regression line is plotted in black.

We compared tuning quality indices in individual trials with smooth pursuit direction errors. The behavioral error was calculated from the difference between mean pursuit direction and individual pursuit directions (Figure 4C). The directions of eye velocity were estimated for individual trials by taking arctan of averaged horizontal and vertical components in a 30 ms time window (across the entire period between 0 to 200 ms from pursuit latency). If the encoding accuracy that we estimated was related to the underlying neural processes that control the directions of pursuit initiation, it would be correlated with behavioral errors. Therefore, we calculated the trial-by-trial correlations between the encoding accuracy estimations and amplitude of the direction errors. Encoding accuracy estimations at the time window between 50 and 100 ms had predictive power over the size of pursuit direction errors after 40 to 50 ms from pursuit latency (approximately 140 to 150 ms from motion onset, Figure 4D, p = 0.007, two-tailed cluster-based permutation test, 5000 permutations). We did not observe a significant correlation between the two when we used the encoding accuracy estimations from other time windows (25 to 75 ms, 75 to 125 ms, 100 to 150 ms, and 125 to 175 ms from motion onset, Supplementary Figure 1). Direction tuning curves at the time window between 50 and 100 ms may have been unique because EEG representations of direction responses were relatively strong, and it occurred before the eyes started to move. Direction tuning responses at earlier times may have been too weak, and responses at later times may have contained the motor behavior or planning components. Therefore, encoding accuracy in the time window of 50 to 100 ms would show how strong the visual motion representation is. This suggested that the trial-by-trial changes in the encoding of *sensory* motion direction had accompanying changes in the direction of motor behavior.

This correlation also suggested that the quality of the direction representation in sensory response was predictive of the reliability of pursuit direction. To investigate this, we divided trials by the quality of encoding. We designated the upper and lower 46% of trials according to encoding accuracy and termed them high and low accuracy trial groups. Our prediction was if the encoding accuracy of the motion direction representation was low, pursuit behavior would be more variable. When we observed the pursuit direction at 50 ms from pursuit latency, the standard deviation of the behavior was larger in the low accuracy trial group than in the high accuracy trial group in an example participant (Figure 4E, variance test, p = 0.024). Across the 15 participants, the standard deviations of pursuit direction at 50 ms from pursuit latency were significantly lower in the high accuracy trial groups than in the low accuracy trial groups (Figure 4F, within-subject design t-test, p = 0.0008).

Finally, we tested if encoding quality could explain the individual differences across participants. Some participants’ motor behaviors were more variable than others, and that may have depended on how well the stimulus motion direction was represented in EEG responses. Therefore, we compared the standard deviation of pursuit directions with the average encoding accuracies across the 15 participants. Figure 4G shows the correlation (Spearman’s rho) between the two measurements across individual participants, as a function of time (from pursuit latency). We found significant negative correlations between 50 and 65 ms, and 145 and 160 ms (p < 0.05). Figure 4H shows the relationship between the encoding accuracy and standard deviation of pursuit direction measured at 50 ms from pursuit latency. This significant negative correlation suggested that the quality of multivariate EEG representation of *sensory* motion direction was predictive not only for within-subject behavioral variation but also for between-subject behavioral variation. We also observed significant correlations across participants when we used the encoding accuracies at later times (75 to 125 ms, 100 to 150 ms, and 125 to 175 ms from motion onset, Supplementary Figure 1D, F, H). Here, the significant correlations started appearing from 50 ms after pursuit latency; therefore, the time windows used for estimating encoding accuracies did not overlap with those for the significant correlations (with the exception of the last one) given that the nominal pursuit latency was approximately 100 ms from motion onset. Although there were significant correlations across participants in these time windows, we did not observe a significant effect of encoding accuracy on behavioral variation across trials (Supplementary Figure 1C, E, G). Therefore, the early, pure sensory direction representation (encoding measured at 50 to 100 ms) appears to have a critical influence over open-loop pursuit direction variation.

## Discussion

We have demonstrated that direction information of sensory and pursuit motion in the smooth pursuit eye movement can be decoded from multivariate EEG activity, as early as 20 ms from motion onset. The early sensory responses in EEG activity patterns are generalized to later direction responses during open-loop smooth pursuit. The encoding accuracy, estimated by the robustness of the direction tuning curves, is predictive of the size of pursuit direction error and variability of pursuit direction within and across subjects. To our knowledge, this is the first demonstration of smooth pursuit direction representation in multivariate EEG activity.

### Open-loop pursuit and cross-temporal generalization

Recent papers have shown the importance of the shape of the cross-temporal generalization map for interpreting how neural representations are dynamically transformed (King and Dehaene, 2014; Stokes, 2015). Our data indicated the initial direction responses in temporal generalization map resembled the ramping pattern, which implies sustained gradual ramping of direction information represented in EEG activity (King and Dehaene, 2014). In the later, closed-loop pursuit period, the generalization map followed a diagonal pattern, indicating that there may be a series of dynamic processing stages. This distinctive pattern can be easily explained by the open-loop versus closed-loop properties of smooth pursuit eye movements. The smooth pursuit eye movements between 0 ms and 100 ms from pursuit initiation, or from approximately 100 ms to 200 ms relative to visual motion onset, are considered open-loop responses (Lisberger and Westbrook, 1985). During this period, eye movements are guided by pure sensory, retinal motion because the feedback signal for correcting eye position has yet to occur in the system. In closed-loop pursuit, the pursuit system constantly corrects eye position so that it can maintain the target at the fovea. Therefore, in the open-loop period, neural representation for motion direction will be maintained, but in the closed-loop period, neural direction representation will dynamically change. Indeed, the cross-temporal generalization map shows that motion direction information is maintained throughout the open-loop period, reflecting sustained ramping sensory motion direction responses in EEG activity, but dynamically changing responses in the closed-loop period due to feedback information for correcting eye position.

### Sensory origin of smooth pursuit variation

In open-loop pursuit, most behavioral variation originates from the variation in estimating sensory parameters of visual motion. Osborne et al. have demonstrated that more than 90% of the trial-by-trial variation in open-loop smooth pursuit eye movements can be explained by errors in estimating direction, speed, and timing of visual motion (Osborne et al., 2005). Hohl et al. have reported that trial-by-trial variation of smooth pursuit speed is correlated with the firing rate of single neural activity in area MT (Hohl et al., 2013). Further, Lee et al. have demonstrated that latency variation of smooth pursuit can be explained by the correlated latency variation between neurons in area MT and multiplicative downstream gain factors (Lee et al., 2016).

According to this view, the correlated neural variation in the sensory area could be the source of behavioral variation. The trial-by-trial covariation of neural activity will effectively decrease the signal-to-noise ratio of population responses (Cohen and Maunsell, 2009; Ni et al., 2018; Zohary et al., 1994), which will result in the deterioration of neural representations of stimulus parameters. Therefore, if the correlated neural variation in the sensory area is the source of behavioral variation, the quality of neural sensory representation of a stimulus will covary with behavioral errors. In our study, we observed a trial-by-trial relationship between the robustness of direction tuning curves during the early, *sensory* response period (50 to 100 ms from motion onset) and the direction error of eye velocity in pursuit initiation (Figure 4D). We believe that the behavioral responses reflecting feedforward processes indeed depends on the *sensory* representation because the trial-by-trial relationship and relationship between encoding accuracy and variation of pursuit direction disappear when we evaluated the encoding accuracy at later times (Supplementary Figure 1C, E, G). If any motor-related EEG representation has the main influence over behavioral variation, the encoding quality of direction at later times should be more strongly correlated with pursuit direction. The fact that the trial-by-trial relationship disappears when the encoding accuracy is evaluated at later times, even if the overall direction tuning responses at later times are superior (potentially due to the combination of sensory and motor responses), suggests that motor-related EEG responses have minimal effects on the variation of pursuit initiation.

### Multivariate EEG representation of direction as a likelihood function in Bayesian inference

The relationship between encoding accuracy of EEG responses to visual motion direction and behavioral variation of open-loop pursuit within and across subjects is easily explained by viewing sensorimotor behavior under the framework of Bayesian inference (Darlington et al., 2018, 2017; Körding and Wolpert, 2004; Yang et al., 2012). In the Bayesian observer model, the true sensory stimulus would be estimated by an ideal observer evaluating a posterior probability distribution, which is formed by the product of a likelihood function and prior probability distribution. In this scheme, a trial-by-trial variation of Bayesian estimates arises from the variation in the likelihood function. Although we did not request human participants to *a priori* select a certain direction, the significant correlation between the encoding accuracy of visual motion direction and behavior is explainable by the relationship between the trial-by-trial variation of the likelihood function and variation in Bayesian direction estimates. As the neural representation of visual motion direction can be evaluated from EEG activity, we have access to the neural representation of the likelihood function. When the neural representation of sensory evidence is weak (when the encoding accuracy is low), the resultant direction of pursuit will be more variable due to higher variation in the likelihood function. When the neural representation of sensory evidence is strong, the trial-by-trial variation in pursuit direction will be small.

### The effect of saccadic eye movements on EEG activity

Previous EEG studies reported that eye movements themselves can have noticeable effects on EEG responses (Dimigen et al., 2009). In particular, EEG gamma activity can be the result of micro- and macro-saccades (Kovach et al., 2011; Yuval-Greenberg et al., 2008). Therefore, the effect of eye movements is a potential concern in this study because eyes move continuously in the smooth pursuit eye movement task. Several of our findings strongly argue against this possibility. In all analyses, we removed trials with any saccadic or micro-saccadic eye movements in the time window between −100 ms and 250 ms from visual motion onset. The exclusion criterion during the fixation (from −100 ms to 80 ms relative to motion onset) was eye speed of 5°/s. During the pursuit (from 80 ms to 250 ms), we discarded any trials that were considered as saccades through visual inspection, with speeds higher than approximately 20°/s (target speed was 16°/s). Given that the effect of saccadic eye movements on ERP responses in the occipital region start appearing when peak eye speed is higher than 22°/s under checkerboard background conditions and the latency of ERP responses is longer than 100 ms (Dimigen et al., 2009), it is unlikely that micro- or macro-saccadic eye movements had any effect on the direction tuning curves during the open-loop pursuit. Based on ERP response latency to saccades, the effect of saccadic eye movements on EEG activity will appear at 350 ms from motion onset. Therefore, any EEG responses from 0 ms to 350 ms relative to visual motion onset should be free from the effect of saccadic eye movements. Further, considering nominal pursuit latency of 100 ms and the earliest significant direction responses occurring at 20 ms after visual motion, direction responses between 20 ms and 120 ms from motion onset should be considered eye movement-free responses.

### Inverted encoding model

To extract visual motion and pursuit motion direction information from the pattern of EEG activity collected from 64-channel active electrodes, we used the inverted encoding model (Brouwer and Heeger, 2011, 2009). There have been several attempts to use the multivariate pattern analysis of EEG to examine population responses to orientation for understanding working memory (Anderson et al., 2014; Myers et al., 2015; Wolff et al., 2017, 2015) and attention (Garcia et al., 2013). However, attempts to understand the multivariate EEG representation of visual motion in sensorimotor behavior are scarce. Multivariate pattern analysis of EEG activity is a good way to understand underlying neural information processes of smooth pursuit eye movements because it is possible to investigate how direction information is represented in whole brain activity and dynamically changing in a relatively fast time scale (approximately 300 ms from motion onset until the end of the open-loop period).

Since this analysis transforms EEG activity in 64-channel space into direction feature space, we exercised caution to prevent any transformation errors from influencing the results. First, we obtained a sufficient number of repetitions for each direction condition from individual subjects. On average, we obtained 190 trials per direction condition and 952 trials per individual subject after removing saccade trials. Second, we used the “leave one condition out” method to prevent overfitting. When we estimated the transformation weight matrix for the inverted encoding model, we used all trials from all other direction conditions excluding the direction in which we wanted to predict the direction tuning curve. Therefore, we estimated the weight matrix using four direction conditions and predicted the direction condition that was not used in weight estimation processes. If the inverted encoding model only discriminates the differences in EEG activity pattern in each direction regardless of specific direction, then the predicted direction tuning responses do not have to represent the target and pursuit direction. Therefore, we were able to ensure that the direction tuning responses in EEG activity truly represented the target and pursuit direction. Third, we used permutation and converted direction tuning curves into standard scores (see *Methods* for details). Therefore, we ruled out any artifactual components that could originate from the unique structure of data; for example, limited numbers of directions for the estimation of direction tuning curves from the inverted encoding model.

## Conclusions

Multivariate pattern analysis of EEG activity has revealed that the neural representation of initial sensory motion was dominant in the open-loop smooth pursuit eye movements, but was not in the closed-loop pursuit. The neural representation measured immediately before the initiation of smooth pursuit had a predictive power over the trial-by-trial variation of motor behavior responding to the identical sensory motion. In addition, the size of the overall variability in pursuit direction was accounted for by the strength of sensory motion representation, which is readily explainable if we understand the smooth pursuit as a result of Bayesian inference.

---

**Supplementary Figure 1:**
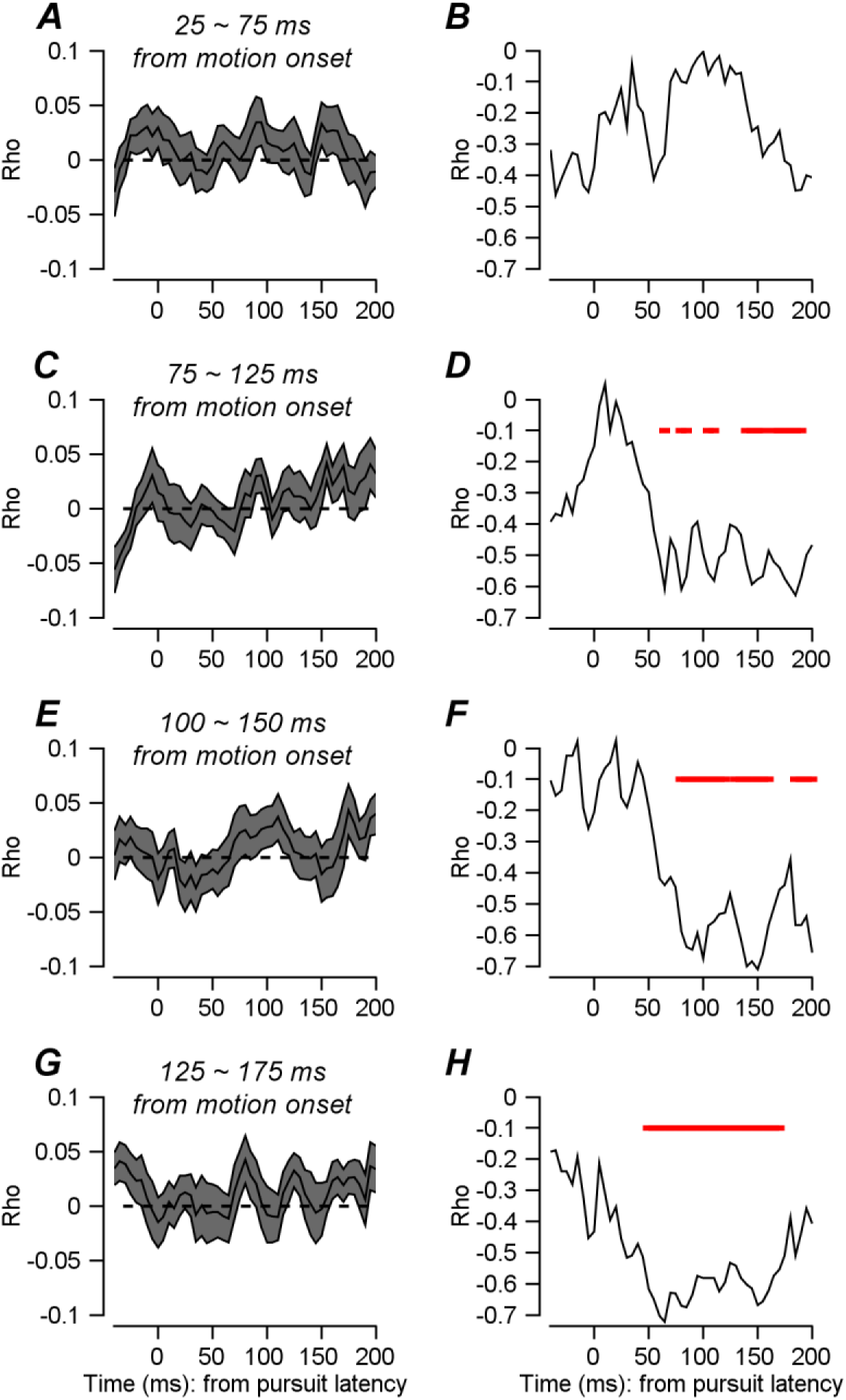
Correlation between encoding accuracy and pursuit behavior. ***A, C, E, G.*** Trial-by-trial correlation between encoding accuracy and pursuit direction errors. Encoding accuracies were estimated at 25 to 75 ms (A), 75 to 125 ms (C), 100 to 150 ms (E), and 125 to 175 ms from motion onset (G). ***B, D, F, H.*** Correlations between average encoding accuracies measured at 25 to 75 ms (B), 75 to 125 ms (D), 100 to 150 ms (F), 125 to 175 ms (H), and standard deviations of pursuit directions across the 15 participants. Standard deviations of pursuit directions are measured from −50 ms to 200 ms relative to pursuit latency. The red bar shows significant correlations (Spearman’s rho, p < 0.05).

## Acknowledgments

We thank Jeongjun Park and Hyomin Yu for assisting with data collection and Dr. Stephen G. Lisberger for constructive comments on earlier versions of this manuscript. This research was supported by IBS-R015-D1. The authors declare no competing financial interests.

Author contributions
Conceptualization, J.L.; Methodology, J.L. and Y.J.K.; Investigation, J.L., W.J., and S.K.; Formal Analysis, J.L.; Writing - Original Draft, J.L.; Writing - Review and Editing, J.L., and Y.J.K.; Funding Acquisition, J.L.

